# Differential modulation of the neurocomputational signature of early and late mismatch responses to spoken words by awareness and task relevance

**DOI:** 10.64898/2026.11.03.757859

**Authors:** Jana Harenbrock, Thomas Straube, Torge Dellert, Insa Schlossmacher

## Abstract

Mismatch responses (MMRs), including the mismatch negativity (MMN) and the P3, are a hallmark of the brain’s capacity to detect unexpected events. They are discussed to be generated by two main processes: adaptation and prediction error (PE). However, MMR research on underlying mechanisms has mostly relied on simple, non-linguistic stimuli such as tones, and it remains unknown how awareness and attentional conditions affect MMRs to auditory verbal stimuli, that, unlike tones, carry lexical-semantic meaning. Here, we conducted a preregistered EEG study that disentangled influences of awareness, general task-relevance of word stimuli, and specific target-relevance of word oddballs on the MMN and P3. In addition, single-trial neurocomputational models were applied to uncover the underlying generative models of the MMRs. Results show that word-related MMN and P3 were only elicited when words were task-relevant, and the P3 increased with target-relevance. Crucially, no word-related MMN emerged under inattention, in contrast to the pre-attentive MMN typically reported for tonal deviants. Neurocomputational modeling revealed that the early part of the task-relevant MMN reflects adaptation processes. PE processes generated later parts of the MMN under task-relevance, and the entire MMN under target-relevance, as well as the P3 independent of relevance conditions. This suggests that the same temporal generation pattern of adaptation and PE underlie mismatch responses to both words and tones, but that, unlike tonal MMRs, eliciting MMRs to words requires the allocation of attention. This attentional dependency may reflect the additional lexical-semantic and multi-level representational demands that spoken words impose on the auditory system.

## 1 Introduction

Navigating through the environment requires the brain to continuously form predictions, detect unexpected and surprising events, and subsequently update beliefs about what happens next. This account has been studied experimentally using electrophysiological components like the mismatch negativity (MMN) and the P3 [1,2]. Experimentally, mismatch responses (MMRs) have been studied in oddball designs, where neural responses to unexpected deviants are compared to responses to frequently presented and thus, expected, standards [1,2]. The MMN can supposedly be observed regardless of whether attention is allocated to the oddball sequence [3–6] or whether participants are unaware of the oddball stimuli [7–10]. Furthermore, there is a line of research using words and pseudowords as oddball stimuli that suggests that lexical processing is pre-attentive [see 11]. In contrast, the P3 is only observed for conscious stimuli and is modulated by their relevance [2,12–14]. When studying awareness and relevance, it is important to note that awareness and task-relevance are often intertwined [15,16], and that task-relevance additionally is nuanced with either presenting the entire oddball sequence or specifically the deviants as targets [for a review, see 17]. Furthermore, past research was often limited to a task-irrelevant condition that was passive or entailed not well controlled distraction (i.e. watching a silent movie, reading a book). Thus, to disentangle these factors, it should be systematically differentiated between conditions in which participants are unaware of the oddball sequence, conditions in which participants are aware of the oddball sequence but the sequence is task-irrelevant, conditions in which the oddball stimuli are task-relevant (e.g., the target is one of the oddball stimuli, but not the deviant) or conditions in which the deviants are target-relevant. Furthermore, it is yet unclear how underlying mechanisms of MMRs are modulated by these conditions.

Previous research proposed that there are two processes, namely adaptation and prediction, that act as complementary components in generating MMRs [17–20]. Importantly, it has been shown that neither process alone is sufficient to explain MMR and that the presence of one process does not rule out the presence of the other [8,21–24]. According to the adaptation framework, the heightened mismatch response can be explained by refractoriness of stimulus-sensitive neural populations towards the standard stimulus. Thus, when a deviant is presented in an oddball experiment, previously non-adapted neurons would respond, which elicits an increased response compared to the smaller response resulting from adapted neurons towards the standard [25–28]. In contrast, the other process proposed to generate MMRs is part of the hierarchical predictive processing account. According to this account, MMRs can be explained by a process where the prediction about an upcoming input is compared to the actual sensory input and whenever a deviant is presented in an oddball experiment (i.e., the actual sensory input contradicts the expectation), a prediction error (PE) would be elicited. This PE would then be propagated upwards in brain hierarchy and subsequently used to generate the next prediction more precisely, enabling efficient information processing [19,29,30]. Predictive processing accounts have also been extended to spoken word recognition, proposing that PEs arise across hierarchically organized levels of representation [e.g., from phonological to lexical and semantic processing; ,31–35]. Additionally, prior research has shown that using words and non-words as stimuli can produce an MMN in different languages [36–38], with previous studies providing initial evidence that the lexical status-MMN (i.e., the deviant was a word among non-word standards) is generated not only by PE but also by adaptation. Specifically, their results show that acoustic properties interact with lexical status in eliciting an MMN and more salient acoustic properties reduce adaptation-related MMN generation [39,40]. However, most studies investigating underlying mechanisms of neural correlates of deviance processing have relied on simple tones as oddball stimuli [for a review, see 17] and the initial finding on word-related MMN is limited to words vs non-words comparisons. Therefore, it remains unclear whether findings of MMN generating mechanisms generalize to more complex, linguistically meaningful stimuli such as different spoken words.

Recently, the idea of a temporal generation pattern of MMRs has been put forward, where both mechanisms occur at distinctive time points. In detail, research points towards a temporal generation of an early MMR, the MMN, where adaptation is a mechanism that is active early, and prediction is the main mechanism to elicit MMRs at later time points [for a review, see 17]. This has been shown especially for the visual modality using control conditions to disentangle the mechanisms [41–44]. While research in audition has been scarce, initial evidence supports the idea of a temporal generation pattern [23,45]. Additionally, results of a recent review suggest that the temporal generation pattern is present in both modalities [17]. In comparison to previously studied auditory stimuli such as simple tones, this temporal generation pattern might be more complex for spoken word stimuli, which additionally carry lexical and semantic information and show a modulated MMN [46–49]. Specifically, results from a first study using word-related MMN at a lexical level point towards a temporal generation pattern when words are deviants among non-word standards [39]. However, the balance between the extent of adaptation and PE depended on acoustic salience and the study is restricted to the lexical level. What remains to be investigated is the temporal pattern of word-related MMN generation including stimuli with lexical-semantic meaning, when trying to control for acoustic salience.

The later occurring MMR, the P3, has been discussed to be generated solely by predictive processes across modalities [50–53]. Therefore, the overall temporal generation pattern for MMRs involves adaptation processes contributing in generating early MMR (specifically, the early part of the MMN), and prediction processes being active at later time points in the MMN and the P3. The extent to which each mechanism contributes to MMR generation is not only affected by the temporal domain, but it has been suggested to be also affected by task-relevance and awareness [8,54–58]. While several studies reported an MMN under task-irrelevant and unaware conditions, only few investigated how this affects the underlying mechanisms directly. Regarding the comparison of task-relevant to task-irrelevant MMN, the findings from previous research show evidence for adaptation and prediction under both relevance conditions [8,54,59,60]. Importantly, only one study found evidence for a modulation by relevance in underlying mechanisms. Specifically, it has been suggested that adaptation predominantly generates the MMN under task-irrelevant conditions, while PE-related processes are more prominent in the task-relevant condition [8]. As mentioned above, task-relevance can be considered more nuanced and should additionally be separated from target-relevance. Regarding the comparison of the generation of the MMN elicited under conditions where participants were aware versus unaware of the oddball sequence, the research to date is limited to vision. This initial research shows evidence for adaptation-related processes generating the MMN under unaware conditions and points towards a transition from adaptation in unaware and task-irrelevant conditions to it being predominantly generated by prediction in the task-relevant condition [8,58]. However, there are no studies in the auditory modality yet.

Therefore, the present pre-registered study (https://osf.io/2v98h/) extends previous studies by using spoken words rather than simple tones as roving oddball stimuli to investigate underlying mechanisms of MMR generation under different awareness and relevance conditions. We aimed to disentangle awareness and relevance by using an inattentional deafness paradigm coupled with different task manipulations, see Figure 1 and Table 1 [61–64]. This allowed us to examine four separate conditions: unaware (participants are unaware of oddball sequence, words are task-irrelevant), aware (participants are aware of oddball sequence, words are task-irrelevant), task-relevant (participants are aware of oddball sequence, words are task-relevant but the deviant is target-irrelevant) and target-relevant (participants are aware of oddball sequence, words are task-relevant and the deviant is target-relevant). Further, we applied neurocomputational modeling to delineate adaptation from PE and categorical models. This allows for the direct comparison of the extent to which each mechanism contributes to MMN and P3 generation depending on the condition. Further, we applied time-resolved neurocomputational modeling to shed light on how the contribution of each mechanism changes within MMN and P3 time windows. This approach should strongly increase our understanding of temporal electrophysiological mismatch responses and how they are affected by awareness, task-and target-relevance.

**Figure 1.**
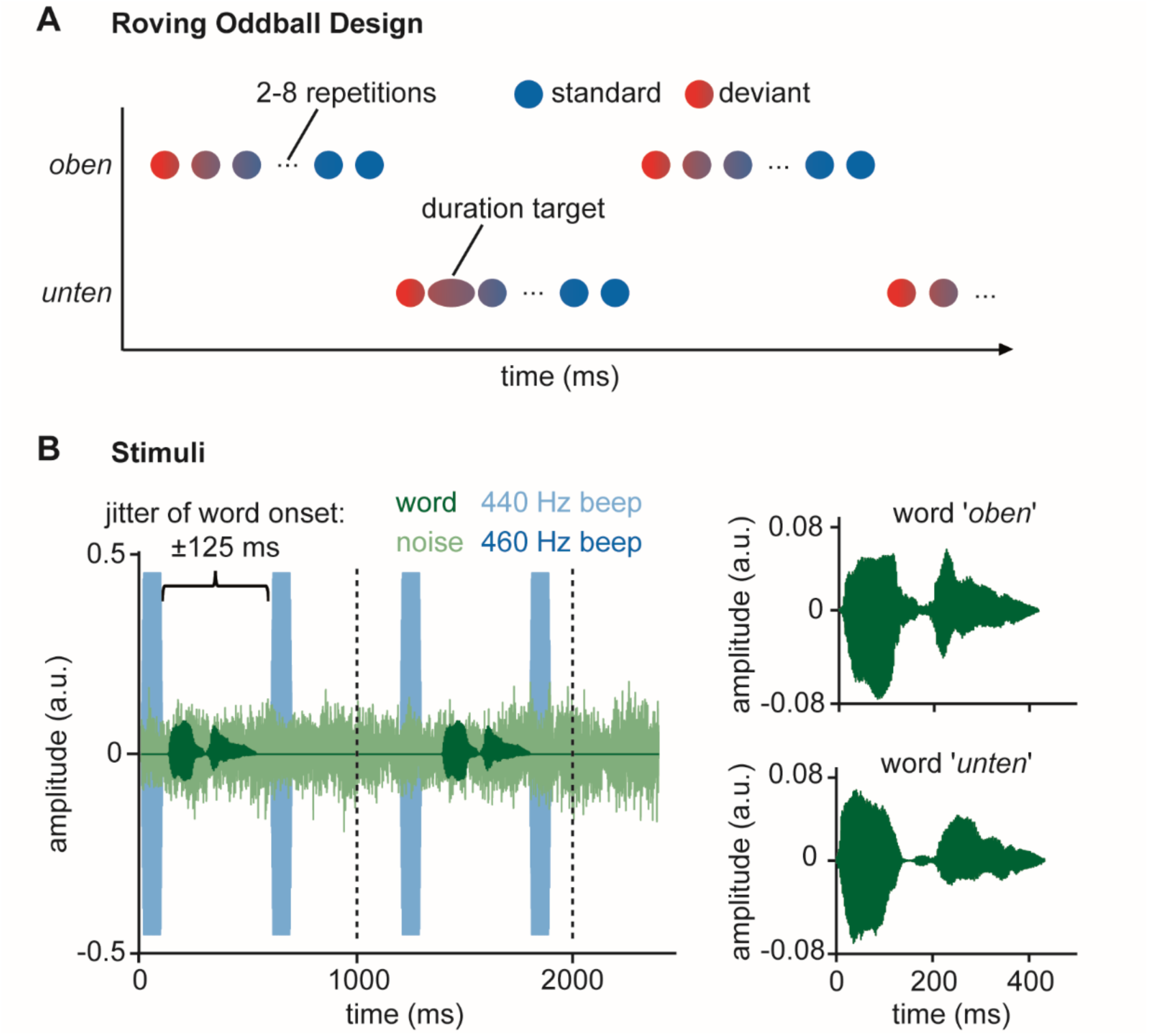
Experimental paradigm and Stimuli. (A) Schematic of roving oddball sequence. (B) Left figure: example trials. One trial consisted of two beeps, continuous noise and a word. Word onset was jittered in relation to the beep onset. The dashed lines separate the illustrated trials. Right figure: Waveforms of the speech stimuli.

**Table 1.** Experimental Conditions.

| Condition |  | Task | Awareness of Words | Task-relevance of words | Target-relevance of deviants |
|---|---|---|---|---|---|
| A | Unaware | Count deviant beeps | No | No | No |
| B | Aware | Count deviant beeps | Yes | No | No |
| C | Task-relevant | Count longer words | Yes | Yes | No |
| D | Target-relevant | Count deviants | Yes | Yes | Yes |

Note. The order of the conditions was pseudo-randomized, with the exception of condition A (unaware) always being presented first.

## 2 Results

### 2.1 Task performance and confidence ratings

Task performance on all tasks was high, indexed by high d’ values (condition A: 4.37, SD = 0.53; condition B: 4.36, SD = 0.46; condition C: 4.41, SD = 0.46; condition D: 4.76, SD = 0.53), indicating that participants were able to comply with the tasks easily. Task performance differed between conditions (*F*(3,126) = 9.08, *p* < .001, BF_incl_ > 100) with holm-corrected post hoc tests showing that participants performed significantly better in condition D as compared to condition A (*t*(42) = −3.69, p_holm_ = .003, BF_10_ = 44.84), condition B (*t*(42) = −4.96, p_holm_ < .001, BF_10_ > 100), and condition C (*t*(42) = −3.56, p_holm_ = .004, BF_10_ = 31.95), Holm-corrected post hoc tests also showed that task performance did not differ between condition A, condition B, and condition C (all p_holm_ = 1, all BF_01_> 3) see Figure 2A.

**Figure 2.**
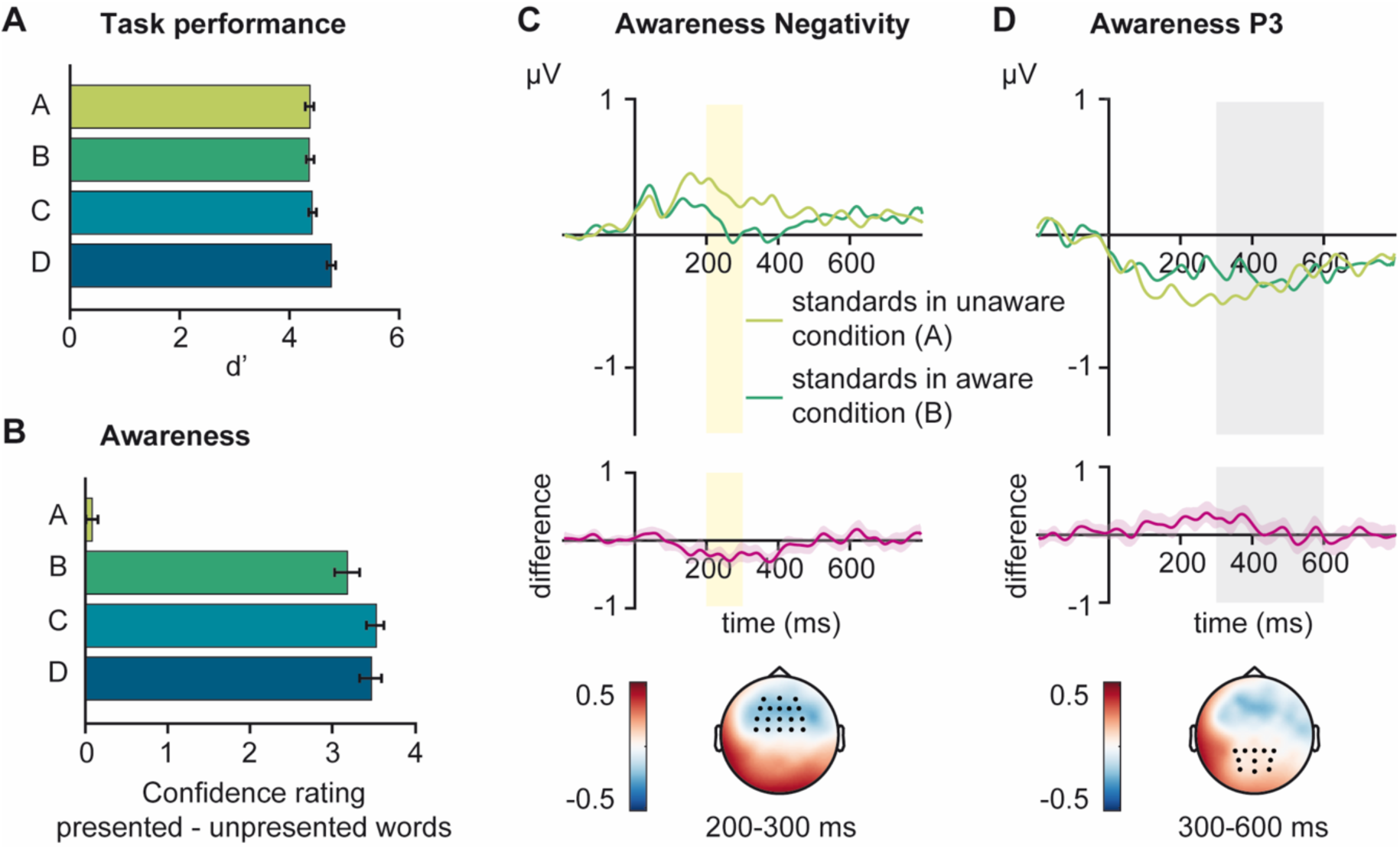
Task performance and awareness manipulation. (A) Task performance measured by d’ for each condition. (B) Awareness measured by rating differences of presented vs not presented words for each condition. (C) Awareness Negativity from 200 to 30 ms (AAN): Waveforms and topography of the significant cluster comparing standards in condition B (aware) to standards in condition A (unaware). The significant cluster comprised the electrodes marked in black and the time interval marked in yellow. Error bars depict standard error of the mean. (D) Awareness P3: Waveforms and topography from 300 to 600 ms (marked in gray) and from electrodes marked in black.

Rating differences (presented – not presented words) were low in condition A, but high in all other conditions (condition A: 0.08, SD = 0.49; condition B: 3.19, SD = 1.02; condition C: 3.53, SD = 0.70; condition D: 3.48, SD = 0.89). Repeated-measures ANOVAs indicated a significant main effect of conditions for the confidence ratings (*F*(3,126) = 225.73, *p* < .001, , *ɛ̂* = .71, BF_incl_ > 100), see Figure 2B. Specifically, holm-corrected post hoc tests revealed that participants were significantly worse in differentiating presented from not presented words in condition A as compared to condition B (*t*(42) = −18.80, p_holm_ < .001, BF_10_ >100), condition C (*t*(42) = −28.83, p_holm_ < .001, BF_10_ >100), and condition D (*t*(42) = −21.33, p_holm_ < .001, BF_10_ >100). Importantly, rating differences in condition A did not significantly differ from zero (*t*(42) = 1.10, *p* = .279, BF_01_ = 3.46), indicating that participants could not differentiate between presented and not presented words. .

### 2.2 EEG

To test the awareness contrast, standards in condition B (aware) were compared to standards in condition A (unaware). A one-sided cluster-based permutation test revealed a significant auditory awareness negativity (AAN) between 200 and 300 ms at fronto-central electrodes (namely, AF3, FC3, F3, C3, FC1, F1, C1, AFz, FCz, Fz, Cz, FC2, F2, C2, AF4, and FC4; cluster mass = −1379.13, *p* < .001, *d* = - 0.54, see Figure 2C). For the P3, no significant cluster was found, and testing the average against zero also revealed no significant effect but also no evidence for the absence of an effect (*t*(42) = 1.33, *p* = .190, *d* = 0.20, BF_01_ = 2.67).

Cluster-based permutation test revealed a significant MMN effect in condition C between 200 ms and 257 ms at fronto-central electrodes (namely, FC3, F3, C3, FC1, F1, C1, FCz, Fz, Cz, FC2, F2, FC4, F4, and AF4; cluster mass = −224.42, *p* = .026, *d* = −0.36), and in condition D between 225 ms and 293 ms at electrodes FC3, F3, AF3, FC1, F1, Fz, F2, F4, AF4 and AFz (cluster mass = −408.16, *p* = .009, *d* = - 0.46), see Figure 2A. Cluster-based permutation tests detected no significant clusters for MMN in conditions A and B, see Figure 2A. To further substantiate absence of MMN in these conditions, the averages over electrodes of interest and intervals of interest were tested against zero. Evidence for the absence of effects was found in condition A (*t*(42) = 1.17, *p* = .25, *d* = 0.18, BF_01_ = 3.22), but not in condition B (*t*(42) = 2.55, *p* = .015, *d* = 0.39, BF_10_ = 2.87) where a significant effect of positive polarity was found. Further, testing cluster averages of the MMN did not reveal any differences between condition C and D (*t*(42) = −1.58, *p* = .123, *d* = −0.24, BF_01_ = 1.94).

Using cluster-based permutation tests, a significant P3 effect was found in condition C and D between 300 and 600 ms at electrodes P3, P1, Pz, P2, P4, PO3, POz, PO4, O1, Oz, and O2 (C: cluster mass = 2267.47, *p* < .001, *d* = 0.56; D: cluster mass = 5365.00, *p* < .001, *d* = 1.13), see Figure 2B. Cluster-based permutation tests detected no significant effect for P3 in conditions A and B, see Figure 2B. To further substantiate the absence of P3 in condition A and B, the averages over electrodes of interest and intervals of interest were tested against zero. Evidence for the absence of effects was found in condition A (*t*(42) = −0.78, *p* = .442, *d* = −0.12, BF_01_ = 4.57), and no significant effect was found in condition B (*t*(42) = −1.49, *p* = .143, *d* = −0.23, BF_01_ = 1.90). Further, testing cluster averages of the P3 revealed a significant difference between condition C and D (*t*(42) = 7.37, *p* < .001, *d* = 1.12, BF_10_ > 100) with a larger MMR in condition D than condition C.

### 2.3 Single-trial Neurocomputational Modeling

Single-trial estimates were computed in the significant MMN and P3 time windows around the individual peak of each participant, separately for the two ERP components and each condition (MMN condition C: M = 227 ms, SD = 16 ms, min = 203 ms, max = 254 ms; MMN condition D: M = 263 ms, SD = 17 ms, min = 231 ms, max = 291 ms; P3 condition C: M = 456 ms, SD = 86 ms, min = 305 ms, max = 596 ms; P3 condition D: M = 492 ms, SD = 75 ms, min = 311 ms, max = 596 ms). For time-resolved neurocomputational modeling, non-overlapping time windows of 10 ms were defined for electrodes and intevals of significant clusters (condition C: from 200-260 ms for the MMN and from 300-600 ms for the P3; condition D: from 220-300 ms for the MMN and from 300-600 ms for the P3).

For single-trial neurocomputational modeling of ERP peaks, in condition C, adaptation models outperformed all other models for the MMN (adaptation: PMP = 0.601, BF = 4.52; low-level PE: PMP = 0.3818, BF = 1.85; categorical: PMP = 0.012, BF = 0.04; high-level PE: PMP = 0.006, BF = 0.02), while low-level PE models outperformed all other models for the P3 (low-level PE: PMP = 0.510, BF = 3.13; adaptation: PMP = 0.443, BF = 2.39; categorical: PMP = 0.028, BF = 0.09; high-level PE: PMP = 0.018, BF = 0.05). In condition D, low-level PE models were best to explain both, the MMN (low-level PE: PMP = 0.565, BF = 3.90; adaptation: PMP = 0.292, BF = 1.24; categorical: PMP = 0.080, BF = 0.26; high-level PE: PMP = 0.063, BF = 0.20) as well as the P3 (low-level PE: PMP = 0.686, BF = 6.56; adaptation: PMP = 0.312, BF = 1.36; categorical: PMP = 0.001, BF = 0.004; high-level PE: PMP = 0.001, BF = 0.002).

Bayesian model comparison of time-resolved EEG data reveals a specific temporal pattern of MMN generation in condition C. We found that adaptation generates the earlier part of the MMN time window of interest from 200 ms until 230 ms. Then, low-level PE models outperform all other models in the later part of the MMN time window from 230 ms until 250 ms, see Figure 5A. There is inconclusive evidence about which models explain the data best in the time window around 260ms. For the P3, the low-level PE model family outperformed all other model families at most time points, except from the time window around 360 ms, where adaptation performed best, and from the time windows around 320 ms, 350 ms, and 550 ms, where inconclusive evidence about the best model was found, see Figure 5C. In condition D, BMC revealed that the MMN was best explained by low-level PE in the time window around 220 ms and from 240 ms until 300 ms. For the time window around 230 ms, inconclusive evidence about the best model was found, see Figure 5B. For the P3, we found that low-level PE was best to explain the entire time window of the P3 from 300 ms until 600 ms, see Figure 5D.

**Figure 3.**
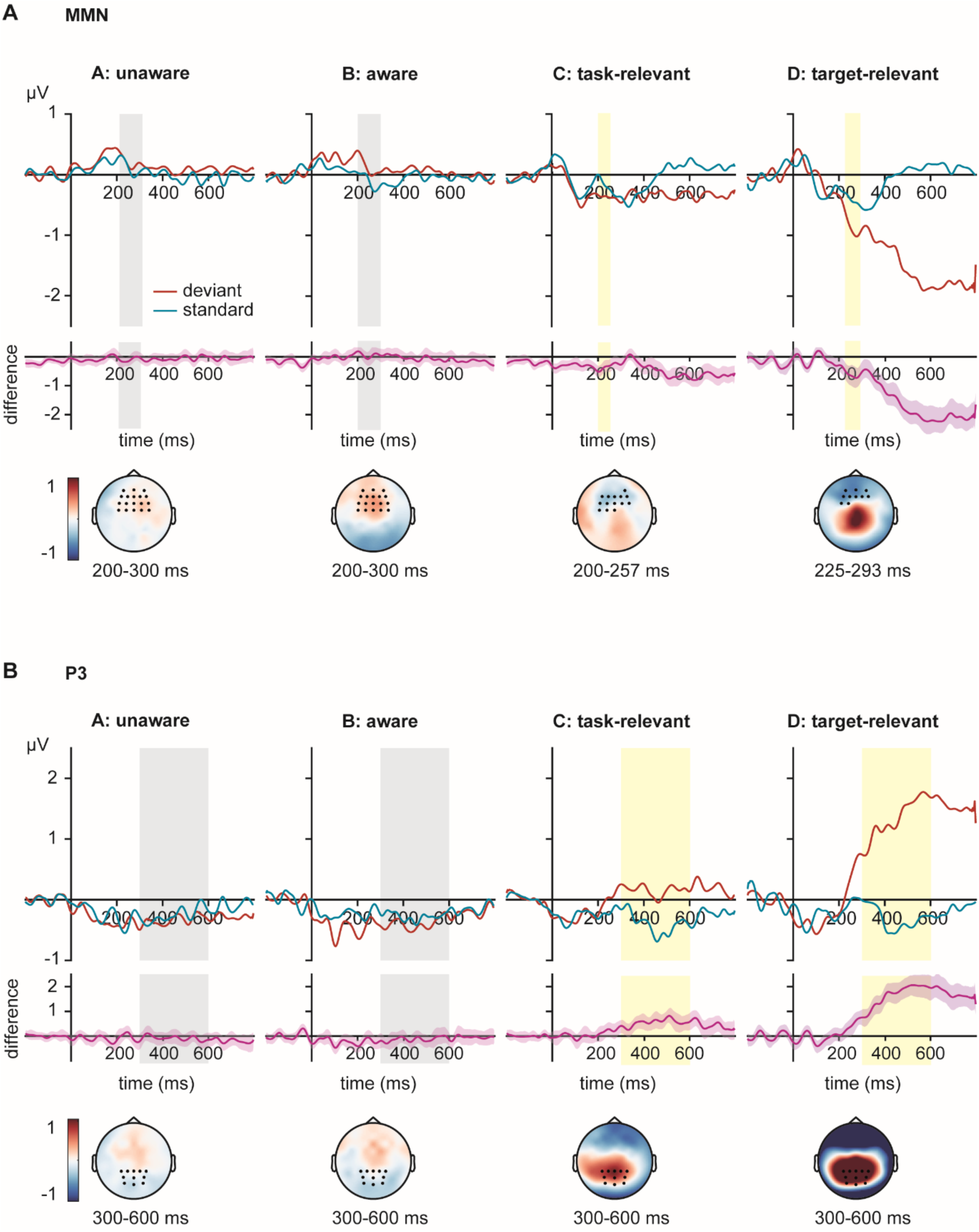
Electrophysiological measures of deviance processing. (A) MMN in all four conditions. The two plots to the left (condition A (unaware) and condition B (aware)) show waveforms and topographies from 200 to 300 ms (marked in gray) and from electrodes marked in black. The two plots to the right (conditions C (task-relevant) and condition D (target-relevant)) depict the significant MMN clusters in the respective conditions. The significant cluster comprised the electrodes marked in black and the time interval marked in yellow. (B) P3 in all four conditions. The two plots to the left (condition A (unaware) and condition B (aware)) show waveforms and topographies from 300 to 600 ms (marked in gray) and from electrodes marked in black. The two plots to the right (conditions C (task-relevant) and condition D (target-relevant)) depict the significant P3 clusters in the respective conditions. The significant cluster comprised the electrodes marked in black and the time interval marked in yellow.

**Figure 4.**
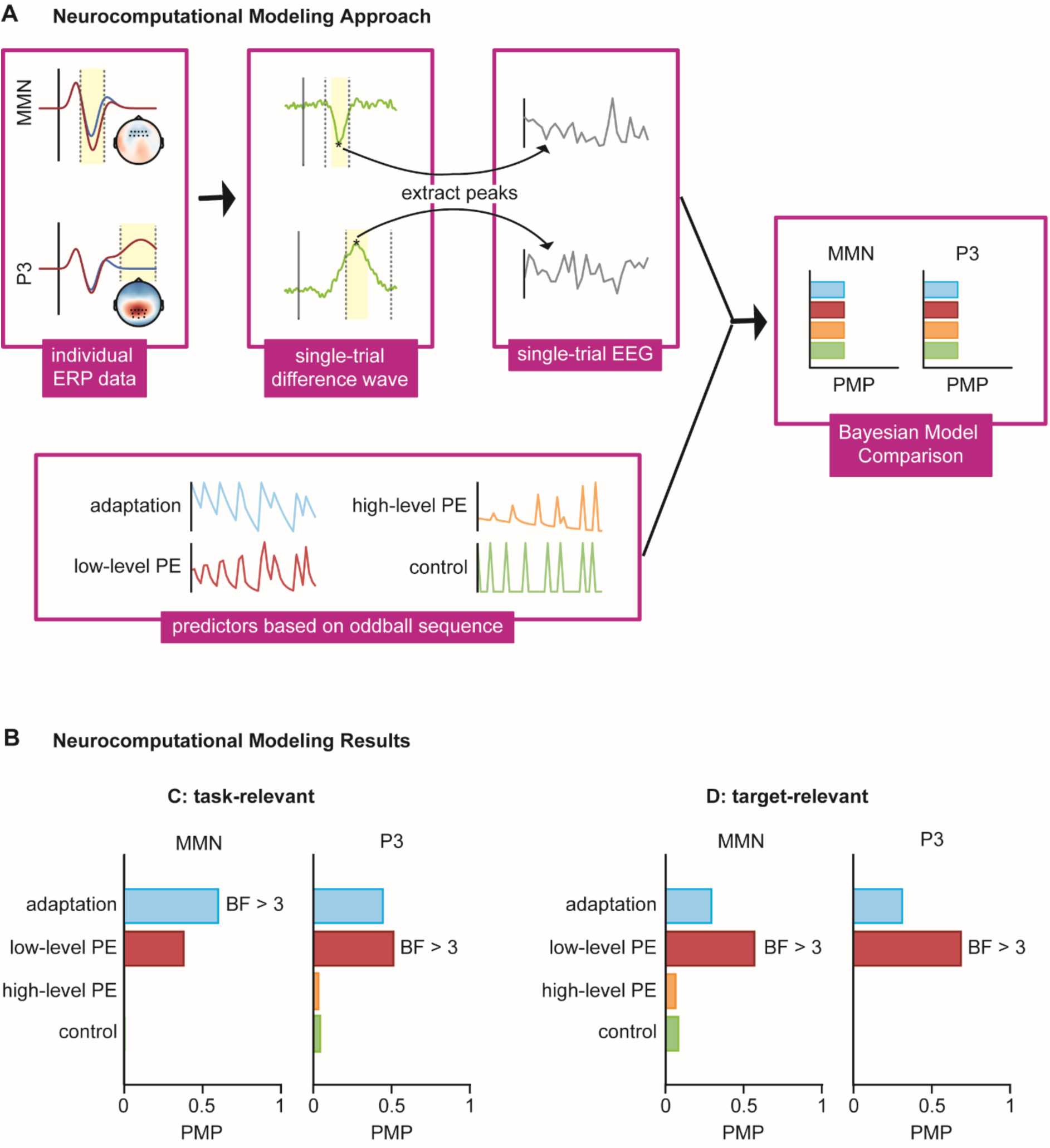
Neurocomputational Modeling. (A) Schematic of the neurocomputational modeling approach. (B) Results from neurocomputational modeling of significant ERPs in condition C (task-relevant, left plots) and condition D (target-relevant, right plots). In both conditions, the left plot shows modeling results for the MMN, and the right plot shows modeling results for the P3. Model evidence is measured by posterior model probabilities (PMP) at the family level and by Bayes Factors (BF) for the best model family.

**Figure 5.**
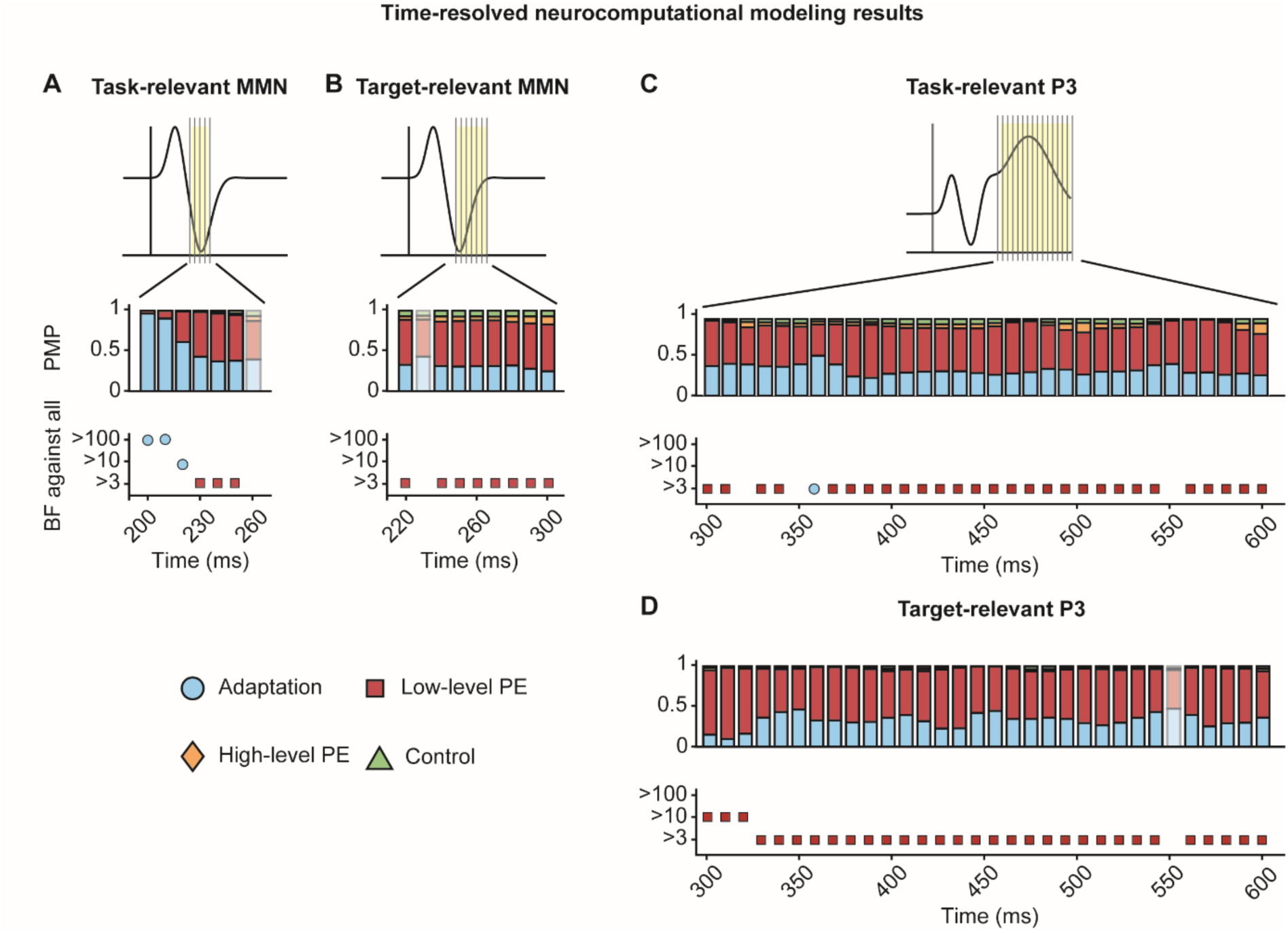
Time-resolved neurocomputational modeling. Upper plots: posterior model probabilities (PMP) at the family level for post-stimulus latencies. Shaded PMP bars correspond to latencies for which no model family dominate substantially (Bayes Factor (BF) < 3). Bottom plots: Relative evidence of each model family against all other families. Only families with a substantial level of evidence (BF > 3) are displayed. Note that we modelled significant time windows, and therefore the modeled latencies of the MMN differ between conditions. (A) time-resolved neurocomputational modeling of the MMN in condition C (task-relevant) from 200 to 260 ms. (B) time-resolved neurocomputational modeling of the MMN in condition D (target-relevant) from 220 to 300 ms. (C) time-resolved neurocomputational modeling of the P3 in condition C (task-relevant) from 300 to 600 ms. (C) time-resolved neurocomputational modeling of the P3 in condition D (target-relevant) from 300 to 600 ms.

## 3 Discussion

In this study, we investigated auditory MMR to spoken words under different awareness and relevance conditions and examined the contributions of adaptation and PE to explain temporal neural signatures of auditory word-related MMR. To disentangle awareness and relevance, we used inattentional deafness and task manipulations to create four conditions: unaware (participants were unaware of the oddball sequence), aware (participants were aware of oddball sequence, but it was task-irrelevant), task-relevant (the oddball sequence was task-relevant, but deviants were target-irrelevant), and target-relevant (deviants were targets). Both MMR components, the MMN and the P3, were only observed when the oddball sequence was task-relevant and target-relevant, but not when it was task-irrelevant (i.e., unattended as in the unaware and aware conditions). Neurocomputational modeling revealed that the MMN to words is mainly driven by adaptation in the task-relevant condition and predominantly generated by PE processes under target-relevance. Further, the word-related P3 is mainly generated by PE in both conditions.

For early MMR, a significant MMN was observed in the task-as well as in the target-relevant condition that did not significantly differ between the two conditions. This is in line with previous studies showing an MMN under the allocation of attention [4,65] and further strengthens the suggestion that MMN is modulated by attention to the entire sequence pattern rather than by attention to the deviants. Specifically, it has been proposed that attention is necessary for formation of standard representation, while the following deviance-detection process is largely independent of attention [5,6]. In contrast to previous studies, we did not observe a significant MMN in the unaware condition [7–10]. However, one study has suggested that at least partial awareness of the standards is needed to elicit an MMN using tones [66]. The inattentional deafness paradigm applied in the present study rendered all oddball stimuli unconscious (see behavioral results and AAN), and therefore, there was no awareness of the standard stimuli that could elicit an MMN. Interestingly, we further did not observe a significant MMN in the aware condition, which would correspond to the typical pre-attentive MMN. Instead, we found a significant effect for a positivity in the typical MMN time window.

One explanation for the absence of an MMN in the unaware and aware conditions might be that the oddball sequence consisted of words. In contrast to tones, which are typically used in oddball paradigms, words have different deviant onsets: the word onset (in the present study the vowel *o* or *u*), the consonant onset (*b* vs *t*), and the onset of the meaning of the words as well as their interpretation (the words *oben* (above) and *unten* (below) are opposites). Hence, using words as stimuli may have increased the variability and attenuated the MMN in the absence of attention, possibly leading to the absence of MMN and its reversed polarity. This interpretation is consistent with psycholinguistic models of spoken word recognition, in which lexical access is not tied to a single acoustic event but unfolds over time as the input narrows multiple candidate words down to a unique match [67,68], and which can be seen in temporally dissociable phonological and semantic mismatch EEG signals [e.g., 69]. Unlike simple tone deviants, words may therefore not elicit one clear mismatch signal but instead lead to an altered mismatch signal based on the multiple levels of word representation [46,70]. The elicitation of an atypical mismatch signal (as seen in the reversal of polarity in the aware condition) or its absence (as seen in the unaware condition) might have additionally been enhanced by the absence of attention and the noise that we added to induce inattentional deafness, which has itself been shown to attenuate the MMN [71–74]. Furthermore, while most auditory oddball studies use a passive paradigm where participants are instructed to focus on a task in a different modality [e.g., watching a movie while listening to the oddball sequence; see 17], in the present study, the distraction task was in the same modality. Because attentional control can limit processing of unattended stimuli in favor of attended stimuli to achieve behavioral goals, the processing of the task-irrelevant oddball sequence could have been reduced [75–83].

Neurocomputational modeling revealed that the word-related MMN is mainly driven by adaptation in the task-relevant condition and predominantly generated by PE processes under target-relevance. The finding of PE processes generating MMN is in line with previous research on attended oddball sequences [17,50–53]. Further, our findings strengthens the hypothesis that PE processes generating the MMN increase with relevance while adaptation processes decrease [8]. In contrast to Schlossmacher et al. [8], however, we do not see this differentiation between the aware and task-relevant condition, but instead for task- vs target-relevance. Further, when looking at results of time-resolved modeling, the contribution of adaptation is limited to an early part of the MMN in the task-relevant condition, while PE processes contribute to generate the MMN later and at all time points under target-relevance. This finding integrates well with research on an auditory temporal generation pattern of MMN from adaptation to prediction [17,23,45]. Importantly, the present neurocomputational results extend the finding of an adaptation- and PE-generated word-related MMN at a lexical level (comparing words to non-words) to spoken word stimuli with lexical-semantic meaning [39,40]. Specifically, the findings suggest that general attention to the auditory word stream (i.e., the task-relevant condition) sufficiently established a stimulus-specific sensory memory trace giving rise to early adaptation-driven MMRs, but was not sufficient to activate a prediction of the upcoming word identity and thus, an PE to deviants. The latter seems to require behavioral relevance of the change in the specific lexical-semantic contrast (i.e., counting the deviants in the target-relevant condition). While the finding of PE generating word-related MMN is consistent with hierarchical predictive processing accounts of spoken word recognition [31,33,34], finding effects of adaptation under the allocation of attention and for more complex stimuli such as words might be surprising. However, a visual study has previously shown adaptation-driven MMN under allocation of attention and word-related MMN at a lexical level has been shown to be driven partly by adaptation [39,40,53]. Further, taken together with the temporal pattern of early parts of the MMN generated by adaptation and later parts by PE, this pattern may be interpretable within the predictive-coding framework [18,19,for reviews, see 84]. According to this framework, adaptation is understood as a signature of PE that has been minimized through either increased precise top-down predictions or through a reduction in the precision afforded to ascending PEs [85–89]. Thus, within this view, increased attention and task demands may modulate the precision-weighting of the PE [90]. The allocation of general attention to the words may enable a precise prediction to establish at the level of the standards’ acoustic and phonological features that then suppresses the associated PE (i.e., adaptation processes), while the lexical-semantic content of the deviant remains only weakly weighted in the task-relevant condition (i.e., the deviant is behaviorally irrelevant). In contrast, under target-relevance, the behavioral importance of the deviant may increase the precision afforded to PEs at a lexical-semantic level, resulting in a strongly weighted PE signal. This interpretation would suggest that the temporal and task-dependent generation pattern of MMN reflects differences in how PEs at different levels in word recognition are weighted as a function of task demands.

Similar to previous research, for late MMR, while evidence for absence of a word-related P3 was found in the aware and unaware conditions, a significant P3 was observed in the task-as well as in the target-relevant condition that significantly differed between the two conditions [2,12–14,91]. Specifically, the P3 amplitude was larger in the target-relevant than in the task-relevant condition. This is in line with research showing that the P3 amplitude increases with task-relevance of the targets [92,93]. Further, this might reflect the increased allocation of attention to the deviants when they are targets that is needed for additional goal-directed processes such as decision making [94].

For the P3, neurocomputational modeling revealed that this late MMR component is mainly generated by PE under task- and target-relevance. The finding of a prediction-generated P3 is in line with previous research on task-relevant P3 [50–53]. This main finding is supported not only by neurocomputational modeling of the ERP peak, but also by time-resolved modeling. While the P3 is generated by PE for the entire time window under target-relevance and for most time points in the task-relevant condition, results in this condition show inconclusive evidence for a few time windows indicating that the models we used might not be sufficient to explain the data at these time points. Further, adaptation was found to be the main generation mechanism at one time point under task-relevance. While this could be true, it is more likely that modeling results of these single non-consecutive time points are affected by high signal-to-noise-ratio, especially when considering that according to posterior model probabilities, PE seems to be a very likely model to explain the data at all time points. Thus, results can be simplified to PE being the main generation mechanism of task- and target-relevant P3 for most of the P3 time window. Interestingly, we did not see a difference for the main generation mechanisms between task- and target-relevant conditions. This is surprising because the significant difference in amplitude between these conditions implied presence of additional goal-directed processes such as decision making [94]. While we did not include models reflecting, e.g., decision making processes, we did include higher-level PE models as well as control models. If such higher-order processes were present for generating the auditory target-relevant P3, we would expect those models to outperform the low-level PE models. In contrast, the present study provides evidence for low-level PE processes generating the P3 under target-relevance. This could point towards a specific processing of the word-related P3, where the use of words involves multiple processing stages such as lexical and semantic meaning [95–97]. According to hierarchical predictive coding accounts of spoken word recognition, PEs are computed separately at these levels of representation with each level contributing distinct temporal signals [31,33,34]. Specifically, the N400 component indexing semantic processing overlaps with the P3 time window and has been linked to PE processing [98–100]. Therefore, the contribution of PE processes and the extent to which it is masked by other processes may differ across stimulus types and for spoken words. We could capture only the earlier PE-related segmental component, while higher-level PE linked to e.g., decision making in the target-relevant condition might occur later and could not be adequately modeled.

To conclude, we observed early and late mismatch responses to words with lexical-semantic meaning under task- and target-relevant conditions, but not in the absence of task-relevance (i.e., only when the words were attended), contrary to the traditional pre-attentive MMN reported for simple tones. Further, the P3 but not the MMN amplitude varied systematically with target-relevance. Importantly, the processes generating these MMRs varied systematically along the temporal domain as well as with target-relevance. This suggests that the same temporal generation pattern that has been established for tonal MMRs also underlie mismatch responses to spoken words, but only if attention is allocated to the stimuli. Specifically, adaptation processes were present for early parts of the MMN under task-relevant conditions (i.e., words were task-relevant but deviants were not targets), while PE processes were found to be the main mechanism generating early parts of the MMN under target-relevance. Later MMRs (i.e., later parts of the MMN as well as the P3) were mainly driven by PE independent of target-relevance. Taken together, these findings suggest that eliciting MMRs to linguistically meaningful auditory stimuli depends critically on task-relevance, potentially reflecting the additional lexical-semantic and multi-level representational demands that words impose relative to simple, non-linguistic stimuli.

## 4 Materials and Methods

### 4.1 Participants

51 participants with normal hearing and no reported history of neurological or psychiatric illness took part in the experiment and were compensated with 12€/h. In agreement with our preregistered analysis, two participants had to be excluded due to noisy data, one participant due to EEG malfunctions and six participants were excluded because they reported to be aware of the background stimuli during the unaware condition. The remaining 43 participants (39 female, 5 male) were aged from 20 to 30 (M = 23.73, SD = 2.73). All participants provided informed consent and the ethics committee of the German Psychological Association (Deutsche Gesellschaft für Psychologie, DGPs) approved the study.

### 4.2 Experimental procedure and stimulus material

Participants listened to auditory stimuli via headphones that consisted of three different components: beep stimuli, babble noise, and speech stimuli. Unawareness of stimuli was achieved by using an inattentional deafness paradigm [see 63], where beeps were presented in the foreground, while speech stimuli (consisting of two words presented in a roving oddball design) were presented embedded in the background noise. The experimental procedure included four conditions: (A) participants were uninformed about the words and focused on the beeps (unaware of oddball sequence, words are task-irrelevant), (B) were informed about the words, but still focused on the foreground task (aware of oddball stimuli, but words are task-irrelevant), (C) focused on targets (i.e., words presented with a longer duration) in the speech stimuli (aware of oddball sequence, words are task-relevant, but the deviant is target-irrelevant), or (D) focused on counting deviants (aware of oddball sequence, words are task-relevant and the deviant is target-relevant), see Table 1.

Speech stimuli consisted of the German words *oben* [ˈoːbn̩ ] and *unten* [ˈʊntn̩ ] (meaning above and below, respectively) that were presented alternately in a roving oddball design and were created using a text-to-speech algorithm in MATLAB (Deng, 2023). The words have about the same spoken duration (*oben*: 419 ms, *unten*: 432 ms), similar normalized logarithmic type frequencies (*oben*: 2.15, *unten*: 1.94; Heister et al., 2011), neutral valence and unambiguous meaning. As the word *unten* is acoustically more salient than *oben* due to the prominent consonant t, the word was low-passed filtered with a cut-off of 1000Hz. Extracted speech stimuli were further cropped at the onset to allow for more precise timing, and a rise/fall time of 10 ms was implemented. On average, every 26 trials (jitter: ±12 trials) a randomly chosen word was presented with double the normal duration (duration target, 23 targets in total). Beep stimuli consisted of sinusoidal tones with rise and fall times of 10 ms, a duration of 100 ms, and frequencies of 440 and 460Hz. On average, every 26 trials (jitter: ±12 trials) the first beep in a trial was presented with a frequency of 460 Hz (beep target, 23 targets in total), while all other beeps were presented at 440 Hz. The babble noise was taken from the freely available signal processing information database [101] accessible via http://spib.linse.ufsc.br/noise.html. The noise was further processed by adding the same stream played backwards to the original. The relative volume of auditory stimuli was set to 0.45/0.25/0.22 (beeps/noise/speech). Beeps were separated by an inter-stimulus interval of 500 ms, while the babble noise was presented continuously. Trials were defined by the presentation of one word and lasted 1 s (jitter: ± 100 ms). Due to the jitter in the word onset, onsets of beep and word stimuli varied from trial to trial. Concurrently to the auditory stream, a white fixation cross was presented on a gray screen, and participants were instructed to fixate at all times [see 63].

In the roving oddball design, stimulus sequences of varying length were presented where the first stimulus was treated as a deviant, while, after several repetitions, it turns into a standard (Lieder et al., 2013; Stefanics et al., 2018). Thus, the oddball comparison involved physically identical stimuli and overall, stimuli were presented an equal amount of times. Speech stimuli were presented in sequences of two (6.67% of sequences), three (13.33%), four (20%), five (20%), six (20%), seven (13.33%) and eight (6.67%) repetitions per word and presentation alternated between the two different words.

The experiment consisted of four conditions with 600 stimuli (∼ 10 min, 120 deviants) per condition, see Table 1. For all four conditions, the stimulus presentation was physically identical, whereas the task differed. In conditions A and B, participants were instructed to count the higher beep tones (460Hz in a stream of 440Hz tones). In conditions C and D, participants were instructed to focus on the speech stimuli presented in the background. In condition A, participants were uninformed about the presence of the speech stimuli. The difficulty of the task was designed to elicit a high percentage of inattentional deafness, so that participants were unaware of the presence of the words. In condition B, the task was held constant, but all participants were informed about the presence of the words and thus, were aware of them. In condition C, awareness was held constant, but participants completed a different task, which directed their attention to the speech stimuli. In this condition, participants were instructed to count the words that had a longer presentation duration, so that the speech stimuli were task-relevant. In condition D, participants again completed a different task, which directed their attention to the deviant stimuli among the words. In this condition, participants were instructed to count the sequence changes (i.e., the deviants) so that oddball stimuli were target-relevant. The order of the conditions was pseudo-randomized, with the exception of the unaware condition being the first condition for every participant. After every 200 trials, there was a break during which participants had to indicate how many targets (i.e., beeps, duration targets or deviants, depending on the current condition) they counted and received performance feedback. Before each condition, a short practice block was presented in order to accustom the participants to the task (in condition A, the difference between beeps was gradually decreased over 3 blocks while only noise stimuli were presented simultaneously; in condition B, speech stimuli were presented additionally and this practice was repeated until participants reported having heard the words; in conditions C and D, the volume of the words was higher in order to facilitate practicing the task). After each condition, a short questionnaire was presented asking whether participants perceived speech stimuli. Then, they were asked to rate four different words (*oben*, *unten*, *groß*, *klein*; the latter meaning big and small, respectively) on how confident they were of having heard the word on a 5-point-scale. The awareness questionnaire and ratings relied on Schlossmacher et al. [63]. Stimulus presentation and response collection were controlled by the software Psychophysics Toolbox [102–104]. Stimuli were presented using an M-audio M-Track 2×2 sound card (M-Audio, Cumberland, RI, USA) and participants listened to the stimuli via headphones (Teufel Aureol Real headphones; Teufel, Berlin, Germany).

### 4.3 Data acquisition and preprocessing

EEG signals were recorded from 64 BioSemi active electrodes using BioSemi’s ActiView software (www.biosemi.com). Electrodes were placed using the 10-20 system conforming with BioSemi electrode caps. Furthermore, vertical and horizontal eye movements were recorded with two electrodes attached above and below the left eye (VEOG) and two electrodes attached to the right and left outer canthi (HEOG). Instead of ground and reference, the BioSemi EEG system uses a CMS/DRL feedback loop with two additional electrodes http://www.biosemi.com/faq/cms&drl.htm). Electrical potentials were recorded with a sampling rate of 512 Hz and impedances were held <20kΩ. A build-in analog anti-aliasing low-pass filter of 104 Hz was applied before digitization.

Preprocessing of the EEG data was performed using the FieldTrip toolbox [105] in MATLAB (version R2019b; MathWorks Inc., Natick, MA; http://www.mathworks.com). Off-line filtering of the continuous data used Butterworth filters with half-power cutoffs. Data was band-stop filtered at approximately 49 –51 Hz (roll-off: −24 dB/octave) and at harmonic frequencies (up to 199 –201 Hz) to minimize line noise. A high-pass filter at 0.1 Hz (roll-off: −12 dB/octave) was applied to remove slow drifts. Then, the EEG signal was segmented into epochs of 200 ms before and 800 ms after stimulus onset. Using an independent component analysis (ICA), horizontal and vertical eye movements were removed from the data. Trials containing muscle artifacts and electrode jumps were manually excluded based on visual inspection and bad channels were interpolated. Data was re-referenced from the CMS/DRL to a common average reference. All trials were baseline-adjusted using the average of a pre-stimulus interval from 200 ms to 0 ms before stimulus onset.

For the oddball contrasts, trials of each subject were averaged separately for deviants and standards. We defined the first stimulus of each roving sequence as a deviant and the last stimulus as a standard to obtain equal amounts of stimuli per condition. This resulted in eight waveforms (deviant/standard in all 4 conditions) per participant. For the awareness contrast, we used an average of all standard stimuli (thus, excluding deviants and last standards) separately for condition A and condition B (contrary to the pre-registered analysis of averaging each phase), resulting in two waveforms for each participant. Last, grand mean waveforms of the averaged data were computed.

### 4.4 Quantification and statistical analysis

#### Task performance

We quantified task performance as *d’* and compared it between conditions. In order to probe conscious perception of speech stimuli, we subtracted confidence ratings for words included in the main experiment (i.e., *oben* and *unten*) from ratings for words not included in the main experiment (*groß* and *klein*, meaning big and small, respectively). Rating scores as well as *d’* were analyzed using repeated-measures ANOVAs and *t*-tests. Whenever sphericity was violated, the Greenhouse-Geisser correction was applied and corrected *p*-values as well as *ɛ̂*-values are reported. These analyses were computed using JASP [Version 0.19.3; ,106]. In addition to parametric tests, we applied Bayesian statistics in order to allow quantifying null effects, with BF_01_ denoting the evidence for the null hypothesis and BF_10_ the evidence for the alternative hypothesis. For Bayesian ANOVAs, we reported the inclusion and exclusion Bayes Factors [107]. We used the conventions from Jeffreys [108] to interpret the results of our Bayesian analyses.

#### EEG

A cluster-based permutation test for within-subjects comparisons was used to investigate early and late mismatch responses [109,110]. Cluster-based permutation tests were performed using the FieldTrip toolbox [105] in MATLAB. In order to enhance its power, we chose a time interval from 200 ms until 300 ms in fronto-central electrodes for the MMN [111]. For the P3, we selected parieto-central electrodes and an interval from 300 to 600 ms [2]. Hypotheses were directional, leading to a one-sided cluster-based permutation test. Clusters were formed by two or more neighboring sensors (in time and space) whenever the *t*-values exceeded the cluster threshold (α = .05). The cluster mass, sum(*t*), was calculated by adding all *t*-values within a cluster. The number of permutations was set to 5000, and the significance value for testing the null hypothesis amounted to α = .05. Prior to the analysis, ERPs were down-sampled to 250 Hz and low-pass filtered at 25 Hz (roll-off: –24 dB/octave) to further enhance statistical power [112]. In order to quantify effect sizes of the significant clusters, we computed Cohen’s *d* for each electrode and time point in the cluster. In addition to pre-registered analyses, averages of electrodes and intervals of interest in conditions where no significant clusters were found, were computed and tested against zero to substantiate the absence of effects. Further, conditions with significant effects were compared to each other. For these analyses, *t*-tests were computed JASP [Version 0.19.3; ,106]. In addition to parametric tests, we applied Bayesian statistics in order to allow quantifying null effects, with BF_01_ denoting the evidence for the null hypothesis and BF_10_ the evidence for the alternative hypothesis. We used the conventions from Jeffreys [108] to interpret the results of our Bayesian analyses.

#### Extracting single-trial EEG data

In order to extract single-trial EEG data, all electrodes included in the significant ERP clusters were averaged and an individual difference wave was computed for each participant, condition, and ERP. In addition to pre-registered analyses, we applied single-trial neurocomputational modeling not only to the MMN, but also to the P3. Therefore, the largest peak of the difference wave (negative for MMN and positive for P3) during the time window of the significant clusters was determined. Last, single-trial estimates (y) were computed as the average amplitude ± 25 ms around peak for MMN and as the average amplitude ± 50 ms around the P3 peak, i.e., the time window amounted to 50 ms and 100 ms for MMN and P3, respectively. All artifact-free trials were used for the single-trial analysis, i.e., in contrast to the ERP analysis, we included all standard trials. In addition to the pre-registered analyses and based on very recent publications and proposals [17,23,113], we extracted time-resolved single-trial EEG data to get a more detailed insight into the temporal generation of MMN and P3. Therefore, we computed mean amplitude differences (deviant – standard) averaged for non-overlapping 10 ms time windows across electrodes and intervals included in the significant clusters.

### 4.5 Single-trial Neurocomputational Modeling

#### Modeling Framework

In order to investigate mechanisms that generate mismatch-responses, we compared four alternative accounts: dynamic adaptation, low and high-level PE and categorical control models. The dynamic low and high-level PE were based on precision weighted Prediction Errors (pwPEs) modeled as predictors from two different hierarchical levels of a Bayesian Learning (BL) model (see Section below for more details). All three dynamic accounts are based on the temporal dependency to previously seen stimuli, a feature that can be explored by dynamic modeling of trial-by-trial fluctuations of mismatch processes. For each of these accounts, we implemented several models that vary in specific values of model parameters. These different variants of the main models can be categorized into the model families of adaptation and Bayesian learning (the latter including 2 model families on different hierarchical levels, see Section below for more details). Additionally, we implemented three different categorical models to serve as controls.

#### Categorical models

As control models, we included three different static models: a constant control model, a categorical oddball model, and a train length change detection model.

#### Constant control model

The constant model predicts a constant response to any stimulus. Thus, it is equivalent to a null model that only includes the intercept.

#### Categorical oddball control model

As a second control model, we included a categorical oddball model that closely mimics the averaging procedure in EEG analysis. This model was always 1 if the stimulus was a deviant, and 0 otherwise:

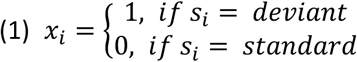

With *s_i_* representing the stimulus type in trial *i*.

#### Train length change detection control model

As a third control model, we included a train length change detection (TLCD) model. This model predicts the magnitude of mismatch to be a response to the deviant depending on the train length of preceding standards:

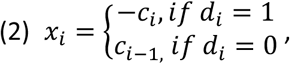

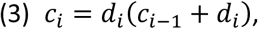

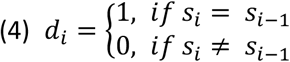

With *c_i_* representing the stimulus train lengths by counting stimulus repetitions, and *d_i_* taking the value 1 whenever the current stimulus is a standard and *d_i_* = 0 resets the current train length to zero [50].

#### Adaptation

We derived adaptation models similar to Lieder et al. [114]. Since we used two different roving words as stimuli, we modelled two different stimulus-selective neuronal populations, *v*_1_ and *v*_2_, resulting in a representation of hidden states by: x_i_ = (x_1,i_, x_2,i_). The responsiveness of each of these neuronal populations was modelled using two exponential processes: Adaptation (*τ*_a_) and recovery (*τ*_r_) [115]. In total, we modeled seven models with different adaptation parameters *τ*_a_ ∊ [3, 10, 20, 30, 50, 100, 200], while *τ*_r_ was always set to be twice *τ*_a_ [23,115]. Neural activity in response to the presented stimuli according to adaptation is thus modeled by:

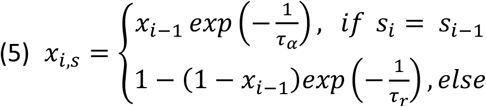

#### Bayesian Learning (BL)

We derived the BL models using the HGF toolbox [8,116–118], which is available at: http://www.translationalneuromodeling.org/tapas. The HGF toolbox enabled us to include generative Bayesian models that are built similarly to Rescorla-Wagner models of reinforcement learning, where the PE is weighted by a learning rate [119]. Additionally, the HGF toolbox includes a hierarchical process with different levels modeled as Gaussian random walks as depicted in Figure 1 [for details, see 116]. Here, we used two different hierarchical levels to model two separate BL model families. For one model family (subsequently referred to as low-level PE models), we used the absolute second level. This corresponds to beliefs about the stimulus property assuming a stable volatility over the time course of the experiment [i.e., the “law” of transition probabilities between stimuli; ,116], that has been proven as relevant in mismatch processing [8,117,118]. We also included models using the parameter of the third level to represent a higher-level PE. These models form a second BL model family (which we will refer to as high-level PE models). To derive pwPE parameters, we used the HGF function tapas_fitModel supplied with configuration files relying on *tapas_bayes_optimal_whatworld_config*, and *tapas_quasinewton_optim_config* (TAPAS release 3.2/HGF version 5.3). Following the implementation of Weber et al. [118] we chose the prior belief about stimulus transitions to be neutral (i.e., both stimuli are equally likely) with µ_2_^(0)^ = 1/2 (var = 0) and a sufficient uncertainty to enable learning with σ_2_^(0)^ = 1 (var = 0; in log-space) and σ_3_^(0)^ = 0.1 (var = 1; in log-space). Because the scale of our third level is arbitrary [120], we chose an initial value of *Κ_2_* = µ_3_^(0)^ = 1 (var = 0). Furthermore, we set ϑ = −1 (var = 2, in log-space) and modeled five models with different Gaussian random walk step sizes between consecutive time points on level 2 with fixed parameters ɷ ∊ [−4, −6, −8, −10, −12], similar to the set of different *τ*_a_ parameters. We extracted trial-wise pwPEs from each of the 5 models at the second and third hierarchical levels, resulting in a total of 10 different trajectories. Absolute pwPEs at the second level were used as regressors for low-level PE models, whereas signed pwPEs at the third level were used for high-level PE models.

#### Model fitting

Model inversion was performed in a first step in each condition in which a significant ERP was found independently for each of the 20 models based on individual ERP peaks (MMN and P3) over all trials. In a second step, model inversion was performed over all EEG trials independently for every model and 10ms time window (to reduce computational burden, we restricted time windows to cover the intervals in which a significant effect was found, ranging approximately from 200 to 250 ms post-stimulus for MMN in condition C, from 220 to 300 and from 300 to 600 ms post-stimulus for P3).

To model mismatch responses, we used a similar approach to previous studies [8,52], where models are estimated by means of a two-level hierarchical general linear model with a random intercept and constant slope on the group level:

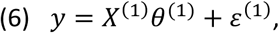

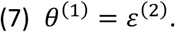

With y representing the concatenated single-trial estimates of all participants, *X*^(1)^ the design matrix, *θ*^(1)^ the regression parameters, and *ε* a random error. The design matrix *X*^(1)^ has *p* + 1 columns and *t* rows, with *p* being the number of participants and *t* being the length of the data vector *y*. *X*^(1)^*_i,j_* is equal to 1 if data *i* is from participant *j*and 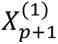 is equal to the predicted values of the tested model (see below). *θ*^(1)^ is of the length *p* + 1 , with the first *p* components representing the random intercepts for each participant *j* and *θ*^(1)^ *p* + 1 representing the regression slope. We specifically wanted to implement a constant slope to ensure that the effects have the ‘right’ direction, e.g., for MMN a negative sign of the slope (in the current study a higher PE value associated with a lower single-trial amplitude consistent with the MMN) and for P3 and fMRI timeseries a positive sign of the slope.

We conducted model inversion using Bayesian linear mixed-effects models as implemented in the *brms* package version 2.23.0 [121,122] using *rstan* version 2.32.7 [123] in *R* version 4.4.1 [124]. The fixed effect for each model regressor was estimated at population level, and subject-specific baseline differences were accounted for by including a random intercept for all subjects. Models were fitted using Markov chain Monte Carlo sampling with four independent Markov chains and a total of 12000 iterations (including 2000 warm up iterations) as recommended in *brms* in order to subsequently compute marginal likelihood using a bridge sampling approach (see Bayesian Model Comparison below). Weakly informative default priors as implemented in *brms* were used with flat priors for regression coefficients, student-*t*(3, µ, 2.5 × σ) distributed priors for intercepts (with µ and σ corresponding to mean and standard deviation of the respective dataset, e.g., single-trial data from MMN in condition C), and student-*t*(3, 0, 2.5 × σ) distributed priors for random effects and residual variance.

#### Model inference

Similarly to Poublan-Couzardot [23], model inference was carried out at the level of model families (namely, categorical models, adaptation models, low- and high-level PE models), aggregating models that share a similar structure but have different parameter values. This BMC procedure allowed us to identify the most plausible mechanism for each ERP (namely, MMN and P3) separately for all conditions revealing significant ERP effects, and additionally for each 10 ms time window at significant time points and electrodes for both ERPs in the respective conditions.

#### Bayesian Model Comparison (BMC)

We used a fixed-effect BMC procedure as implemented in *brms*, following recommendations for physiological mechanisms that are unlikely to vary across subjects such as the MMN [23,125]. Using brms, we computed posterior model probabilities of fitted models from marginal likelihoods that were calculated using a bridge-sampling approach [121,122,126,127]. To compare models (and model families), we used posterior model probabilities (see Formula 9) as well as Bayes Factors (see Formula 10) for a model (family) as implemented by Poublan-Couzardot et al. [23] in the context of a uniform distribution over model space.

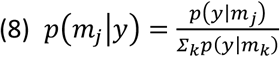

With *p(m_j_|y)* representing the posterior model probability of one specific model *m_j_* out of all models *m_k_* given the observed data *y* and *p(y|m)* representing the model evidence.

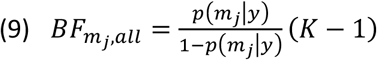

With K representing the number of compared models, all with equal prior probability *p*(*m_k_*) = 1/*K*.

## 5 Acknowledgement

This research was funded by the DFG. The authors declare no competing financial interests. We thank Jennifer Eske, Anja Wiegand, Emily Rupprecht and Larissa Stähler for their assistance in data collection and data preprocessing.

